# The ages of zone of proximal development for retrospective time assessment and anticipation of time event

**DOI:** 10.1101/2020.12.10.419218

**Authors:** G.V. Portnova, A. B. Rebreikina, O.V. Martynova

## Abstract

We aimed to investigate the ability of children aged 5–14 years old (preschoolers, primary schoolers, and preteens) to assess and anticipate time intervals. 287 Russian children aged 5–14 years old and 26 adults of control group participated in our study. The neuropsychological assessment, Wechsler Intelligence Scale for Children and a battery of time-related tests were applied. All groups of children overestimated the event’s duration, although the accuracy of the second estimations increased among the participants aged 6–8 years after a prompt was offered. A zone of proximal development for time anticipation task was detected for children aged 9-11 years, when the prompt could significantly improve the accuracy of time perception. The participants overestimated the duration of both upcoming and past events, with the degree of overestimation being found to be negatively correlated with age. Further, a higher degree of accuracy in terms of time estimation was found to be correlated with higher scores on the attention and memory tests, and accuracy of time anticipation was associated with scores of praxis test.

## I. Introduction

Well known, that the understanding of time is comprised of different aspects, which includes time estimation, subjective time awareness and subjective time perspective (Lehmann, 1967). The ability to estimate time is universal, which means that even infants have a primitive “sense” of time, are able to learn the temporal intervals between two events, and are able to comprehend the durations of events (Brannon, Roussel, Meck, & Woldorff, 2004; Pouthas, Droit, & Jacquet, 1993). At the same time the objective time perception is socially dependent, needs to be internalized and mastered and hence develop throughout childhood (Block, Zakay, & Hancock, 1999). The time judgment does not develop simultaneously and a first progressive increase in the functions associated with time perception has been detected between the ages of three and ten years (Brannon et al., 2007). Children aged 3–5 years could accurately judge the lengths of temporal intervals when they repeatedly experienced the durations of events or the durations of daily activities (Friedman, 1990) and were able to name the seasons (Sharman, Cross, & Vennis, 2004). This developmental transition from making implicit to explicit time judgments begins between the ages of three and six years (Lowe, Beasty, & Bentall, 1983), and by about the age of seven children develop explicit time knowledge (Levin & Zakay, 1989). Yet, prior to the age of ten years, most children do not spontaneously apply explicit time-related strategies (Pouthas, Droit, Jacquet, & Wearden, 1990). Further, the correct judgments concerning durations of time require the sophisticated reasoning abilities that emerge at approximately eight years of age (Piaget, 1981) and accurate temporal judgments being made prior to the age of eight (e.g., Friedman, 1990; Levin, 1982, 1992; Montangero, 1977; Richie & Bickhard, 1988).

There is currently no consensus as to the age at which children must be able to estimate time using units of time (seconds, minutes, hours, days, and years). Moreover, some studies have shown that the ability to estimate the time using a clock, especially a blind clock, is culturally dependent (Kaplan, 2016). For example, there are units of time that are uncommon or even obsolete in relation to European cultures that are widely used in other cultures or communities, such as the Saros, Kalpa, indiction cycle, etc. (Meimaris, 1992). Otherwise, knowledge regarding the various units of time is necessary for explicit time perception. This includes the estimation of a discrete duration of time so as to compare it with a previously memorized standard, which requires the storage and retrieval of long-term memory as well as a comparison with other elements of the working memory (Ciullo, 2015). This ability requires cognitive resources, and it is thus dependent on an individual’s intelligence quotient (IQ), attention level, motivation, and other cognitive abilities. Finally, one of the most complicated processes related to explicit time perception concerns the ability to anticipate the durations of upcoming events. The ability to plan the duration of a given activity develops during maturation, and it requires knowledge about units of time (seconds, minutes, days, etc.), the capacity to recall the durations of previous events, the ability to discriminate between time intervals, and the ability to plan one’s own activity.

At the same time, each cultural dependent or independent ability to perceive time had its own zone of proximal development which could be defined as “the distance between the actual developmental level as determined by independent problem solving and the level of potential development as determined through problem-solving under adult guidance, or in collaboration with more capable peers” (Vygotsky, 1978). So, regarding to development of explicit time perception, the most promising for research seemed to be the investigation of zone of proximal development, which less depended on culture-related abilities to estimate time. We hypothesized that the each time perception ability has similar developmental stages as other cognitive functions, in particular the zone of proximal development (Chaiklin, 2003; Bruner, 1984). In the present study, we aimed to investigate the ability of preschool and school-aged children to assess and reproduce time intervals, and to plan the durations of upcoming events. We attempted to investigate the zone of proximal development (Berk & Winsler, 1995) regarding the ability of children to estimate the time interval using prompt including duration of previous event.

## II. Methods

### A. Participants

The 287 Russian children aged 5–14 years old and 26 adults participated in our study. Participation in the study was entirely voluntary. The participants had no history of brain injury or any psychiatric disorders. Descriptive statistics concerning the participants are presented in Table 1. The participants among children represented four age groups: 5–6 years (preschool), 7–9 years (primary school), 10–12 years (preteen) and 13-14 age (teens).

**Table 1.** Descriptive statistics The results of time perception.

|  |  |  |  |  | Before Prompt |  | After Prompt |  |
| --- | --- | --- | --- | --- | --- | --- | --- | --- |
|  | age | n | Wechsler<br>test value<br>Full-Scale IQ | gender | Schulte<br>time<br>reading | Schulte<br>time<br>assessment | Schulte<br>time<br>reading | Schulte<br>time<br>assessment |
| preschoolers | 5 | 36 | 107±8.3 | 40 girls,<br>41 boys | 143,2±14,5 | 755,7±16,3 | 156,7±19,2 | 762,5±14,7 |
|  | 6 | 45 |  |  | 102,8±13,5 | 483,6±19,2 | 102,2±13,1 | 188,4±15,9 |
| schoolers | 7 | 26 | 109±10.1 | 34 girls,<br>40 boys | 83,2±16,6 | 415,1±20,1 | 79,2±14,3 | 89,5±12,8 |
|  | 8 | 31 |  |  | 74,2±13,9 | 331,7±19,7 | 68,5±12,8 | 79,5±11,2 |
|  | 9 | 20 |  |  | 63,5±12,4 | 89,5±11,5 | 57,8±10,2 | 71,56±10,1 |
| preteens | 10 | 26 | 110±6.9 | 37 girls,<br>41 boys | 46,6±10,4 | 66,8±12,4 | 45,8±9,7 | 54,2±12,4 |
|  | 11 | 26 |  |  | 45,7±11,3 | 60,7±10,4 | 46,3±7,6 | 62,1±13,5 |
|  | 12 | 26 |  |  | 43,1±12,1 | 58,3±13,1 | 45,8±9,9 | 59,3±12,1 |
| teens | 13 | 29 | 111±7.5 | 28 girls,<br>23 boys | 43,4±7,8 | 51,5±9,4 | 39,8±9,5 | 48,7±8,6 |
|  | 14 | 22 |  |  | 42±3,7 | 45,6±8,4 | 37,3±5,4 | 39,3±12,4 |
| adults | 22.4±4.9 | 26 |  | 15<br>women,<br>11 men | 35,1±5,4 | 36,4±7,5 | 32,9±4,9 | 33,3±11,6 |

The participants were recruited from among the children who attend the Center for practical psychology “Equalize”. All data were collected between 2015 and 2018. Eight participants were excluded from the analysis due to data being missing from their test results. A parent or guardian of each participant signed an informed consent form (the protocol in accordance with the Declaration of Helsinki) written in the participant’s native language prior to the tests being administered. The study was approved by the Ethics Committee of the Institute of Higher Nervous Activity and Neurophysiology of the Russian Academy of Science.

### B. Cognitive assessment

Luria’s neuropsychological assessment was also applied in the present study. This involved the assessment of the participants’ visual and verbal memory, time perception, categorization, and naming ability.

The neuropsychological testing included:

1. Quasi-spatial tasks: the first series of tasks included 6 pairs of objects of a barrel and a box with different positions (right, left, behind and etc.). The children were instructed to show a picture with objects in correct position. The second series of tasks included verbally assigned tasks (6 tasks) for spatial or temporal relationships of objects. The number of correct answers was estimated
2. Attention tasks (Schulte tables, see paragraph “Assessment of time perception”)
3. Visual memory tasks: included a set of 5 figures which presented to children for 10-15 seconds with instruction to remember them, further the set was hidden and children should draw figures he reminded. If children couldn’t correctly remind all figures the procedure (with the same figures) repeated again. The number of correctly reproduced figures after the first presentation, the number of presentations required for correct reproduction, and the type of errors (violation of order, distortion of the figures’ elements and etc.) were evaluated.
4. Auditory memory tasks: included verbally assigned 5 words with instruction to remember them and then children should remind words in correct order. The number of correctly reproduced words after the first presentation, the number of presentations required for correct reproduction, and the type of errors (violation of order, contaminations, confabulations and etc.) were evaluated.
5. Coping task: the image of a house with fence and a tree were presented and children were instructed to copy it. The drawing strategy, the number and type of errors (spatial errors, the presence of unnecessary or missing details) were evaluated.
6. Dynamic praxis: neuropsychologist showed the sequence of movements “palm-fist-rib” and (or) “fist-palm-rib” then asks to repeat them. Both hands were examined. The number of repetitions for correct reproduction and type of errors (perseverations, simplification of the program, spatial errors, and stereotypes) were evaluated
7. The choice reaction. The children were instructed to raise his finger in response to a neuropsychologist’s raised fist, and raise a fist in response to a raised finger. First, the experimenter 3 times shows a successive alternation of movements - “finger-fist” (creating a motor stereotype), after that the same movement is presented twice in a row, and then another. The number of repetitions necessary for mastering the task and the number of impulsive errors were evaluated.

Wechsler Intelligence Scale for Children-Fourth Edition (WISC-IV) (Wechsler, 2003) was used in this study to measure the intellectual ability of the participants (Table 1).

No significant differences were found between the groups in this regard. Any participants who scored under 70 on the WISC-IV were excluded from the study.

### C. Assessment of time perception

#### Estimation of event’s duration

The participants were asked to locate the digits on a Schulte table (a 5 x 5 table featuring randomly distributed numbers was used) in ascending order as quickly as possible. After finishing the task, the participants were asked to estimate how much time had passed while they were completing the task (the time taken for the task had previously been determined). After providing their estimations, the participants were told the actual length of time required for the task. This procedure was then repeated for a second time.

#### Anticipation of event’s duration

children were asked to complete a task (contouring a figure). Before they began the task, the participants were asked how long they thought it would take (i.e., to estimate how much time would be needed for the task). After finishing, the participants were asked to assess the duration of the task, that is, to estimate how much time had passed while they were contouring the picture. After providing their estimations, the participants were told the actual length of time required for the task. This procedure was then repeated for a second time with the other picture.

### D. Procedure

All the tests used in this study were administered individually in a quiet and comfortable atmosphere by the author and a psychologist trained in neuropsychological assessment. The testing took approximately 90–120 minutes to complete. To avoid order effects, the sequence in which the tests were administrated was randomly varied (whenever possible) from participant to participant. The participants’ parents were debriefed with regard to the purpose of the study, and they were offered the opportunity to discuss their children’s results with a qualified coinvestigator.

### E. Scoring

The results of neuropsychological assessment were evaluated due to accept previously protocols (Khomskaya, 1987; Luria, 1980).

We measured the participants’ error of estimation event’s duration between the two series of Schulte tables’ reading (accuracy of the time assessment) as |estimated time-real time| and accuracy of error of anticipation of event’s duration as |anticipated time-real time|

## III. Results

### A. Correlations with age, gender and IQ

Age was found to be negatively correlated with the duration of the Schulte task (r < −0.57), the total number of trials required during the verbal memory test (r = 0.44), and the number of missing details during the coping test (r < −0.51). No significant differences were found between the three groups in terms of the WISC-IV scores (Wechsler, 2003). There was no gender differences in time estimation.

### B. Estimation of event’s duration

The majority of participants overestimated the time spent reading the Schulte table. The estimation of Schulte tables’ reading negatively correlated with age (r=-0.68, p=0.0001). Children 5-8 years significantly overestimated events compared to 9 year old and older children that accurately estimated the events (see Figure 2).

After the second attempt at the task, the time estimations became significantly more accurate in the children 6–8 years old (F(10, 302)=29,577, p=0,0000).

The error of the time assessment inversely correlated with age (r = −0.42, p = 0.033) and the results of the visual memory test (the number of correctly recalled figures; r = −0.64, p = 0.004) and directly correlated with the time spent reading the Schulte table (r = 0.59, p = 0.01), the number of mistakes made during the verbal and visual memory tests (order mistakes), and the number of missing details when the participants were asked to copy a picture (r = 0.48, p < 0.05). The participants’ accuracy correlated with the amount of superfluous detail given during the coping task and the number of mistakes made during the quasi-spatial solving task. More specifically, the higher was accuracy the lower was the number of spatial and quasi-spatial verbal mistakes made (r = 0.46, p=0.03).

### C. Anticipation of event’s duration

All the participants overestimated the duration of both the upcoming event and the past event. Further, significant differences were found between the groups (Figure 3). The older children showed lower error of anticipation (r = −0.58, p = 0.008). Moreover, the error of anticipation was found to be positively correlated with the number of mistakes made during the testing of dynamic praxis (r = 0.8, p = 0.001).

|  | Before Prompt |  |  | After Prompt |  |  |
| --- | --- | --- | --- | --- | --- | --- |
|  | contur_plan | contur_real | contur_assess | contur_plan | contur_real | contur_assess |
| preschoolers | 826.3±91,8 | 128,2±25,1 | 815,9±34,5 | 819,4±37,1 | 106,7±15,1 | 812,5±31,4 |
|  | 803,5±85,7 | 118,7±25,4 | 577,3±25,6 | 801,7±28,5 | 119,4±12,4 | 576,9±26,9 |
| schoolers | 656,2±87,2 | 119,2±21,2 | 441,8±31,2 | 614,9±27,5 | 133,7±15,5 | 423,2±37,8 |
|  | 605,3±89,7 | 112,1±14,5 | 310,8±27,5 | 578,6±29,7 | 113,2±14,0 | 349,1±26,9 |
|  | 224,7±41,4 | 124,3±19,4 | 195,1±20,2 | 217,5±21,5 | 126,1±19,8 | 201,5±27,7 |
| preteens | 219,2±42,1 | 84,2±9,6 | 141,2±16,6 | 163,2±24,9 | 91,4±11,1 | 136,4±17,7 |
|  | 205,1±39,4 | 81,5±12,0 | 149,1±17,8 | 172,1±23,2 | 88,7±11,1 | 157,6±12,6 |
|  | 175,2±35,9 | 83,5±11,7 | 115,2±12,8 | 103,8±19,3 | 84,9±13,3 | 85,8±14,4 |
| teens | 121,7±42,4 | 75,6±8,8 | 104,6±12,1 | 104,8±16,1 | 80,2±10,7 | 94,9±9,5 |
|  | 117,7±23,1 | 75,6±4,9 | 99,3±10,2 | 100,4±15,7 | 80,4±9,5 | 91,7±7,8 |
| adults | 98,25±22,5 | 62,1±3,4 | 64,1±4,9 | 85,4±12,8 | 68,8±5,1 | 71,6±5,0 |

**Figure 1.**
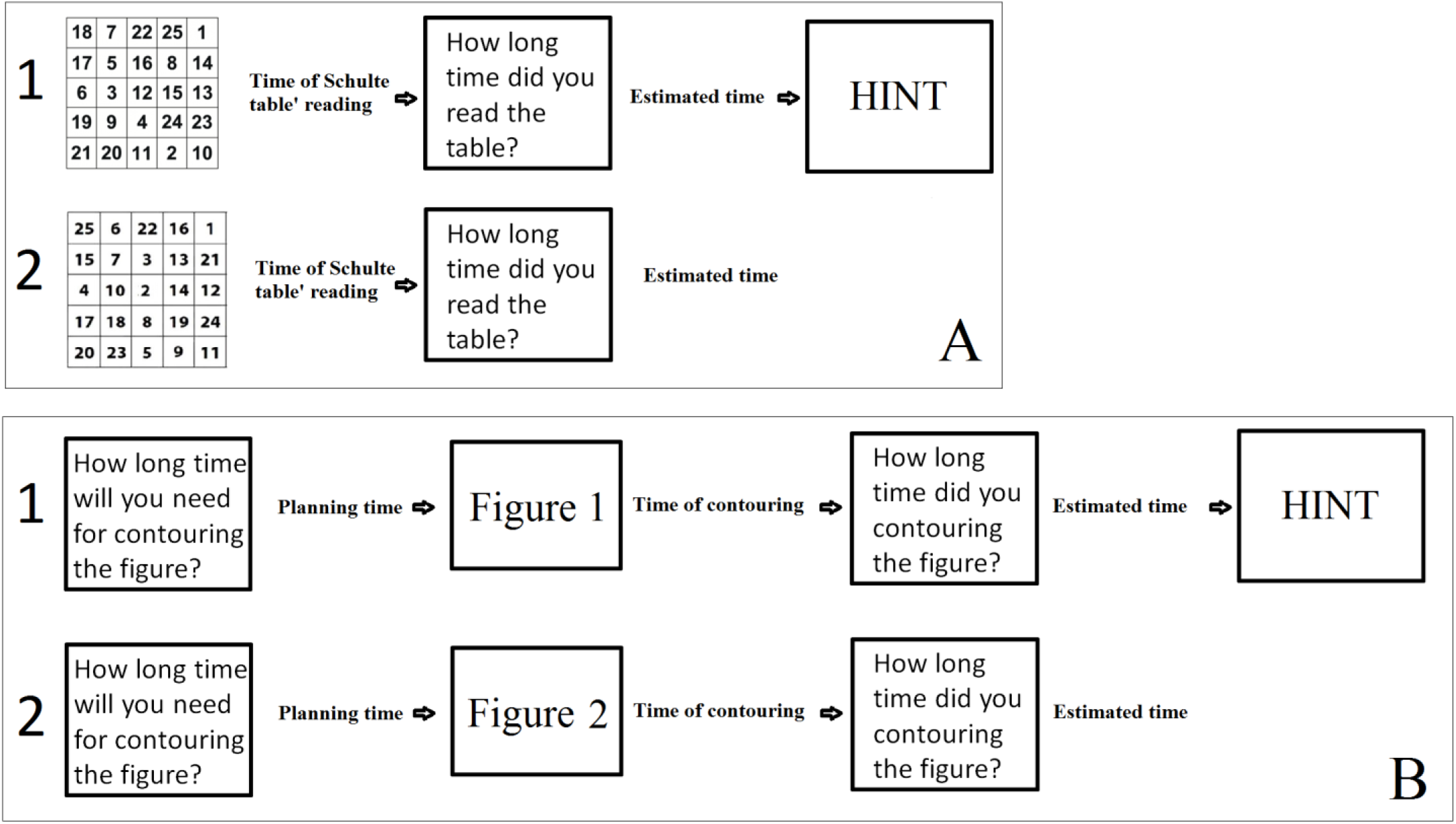
Block-schema of the of the experimental design: A: the task of the event’s duration (Schulte tables’ reading) estimation; B: task of the anticipation of the event’s duration, 1 – first attempt, 2 – second attempt after the prompt. Prompt means, that the participants were told the actual length of time required for the task

**Figure 3.**
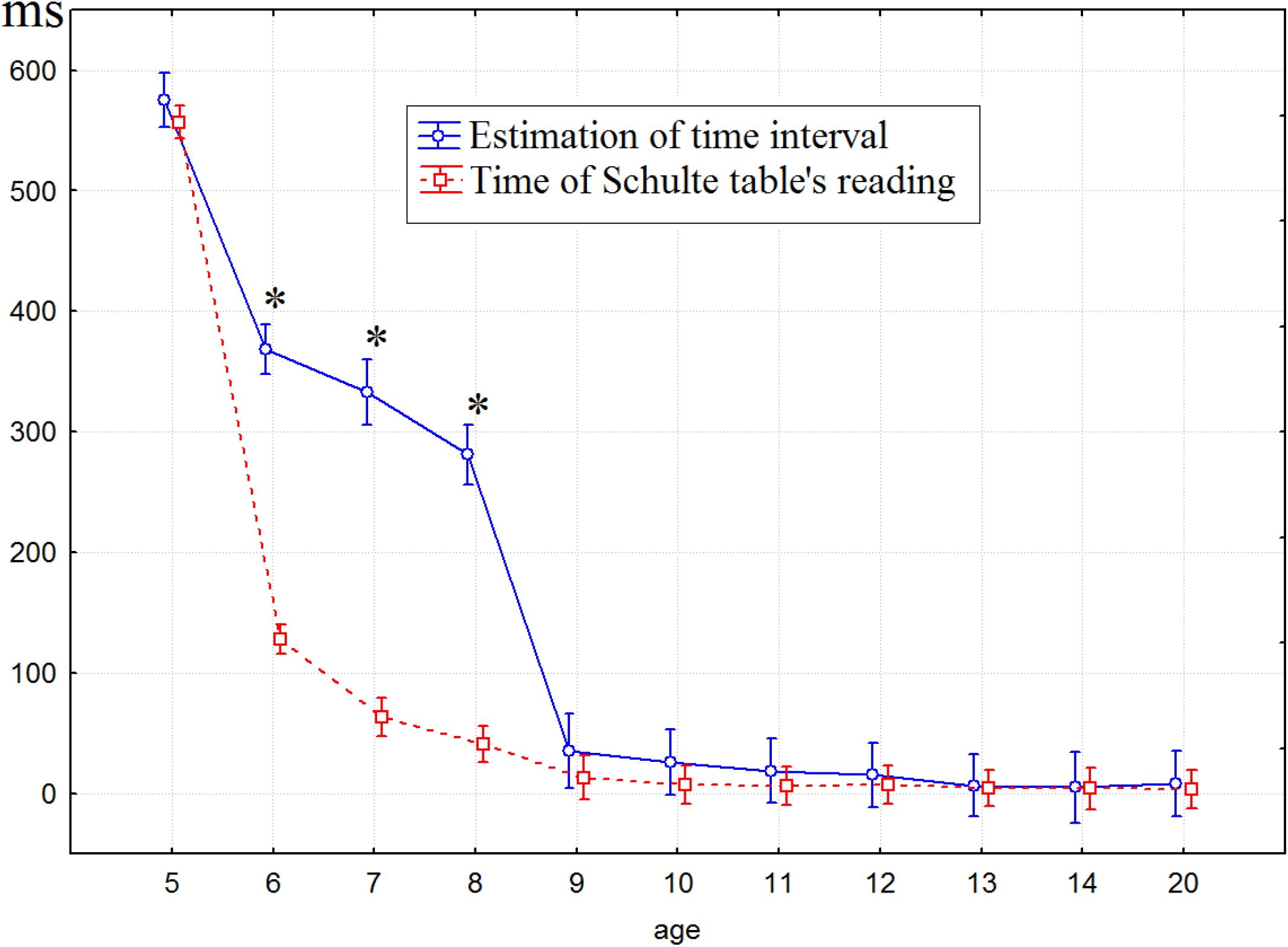
The difference between assessment and real time of tables’ reading (|assessment – real time|). Gray color – first probe, black – second probe. Stars – significant differences between two probes.

**Figure 3.**
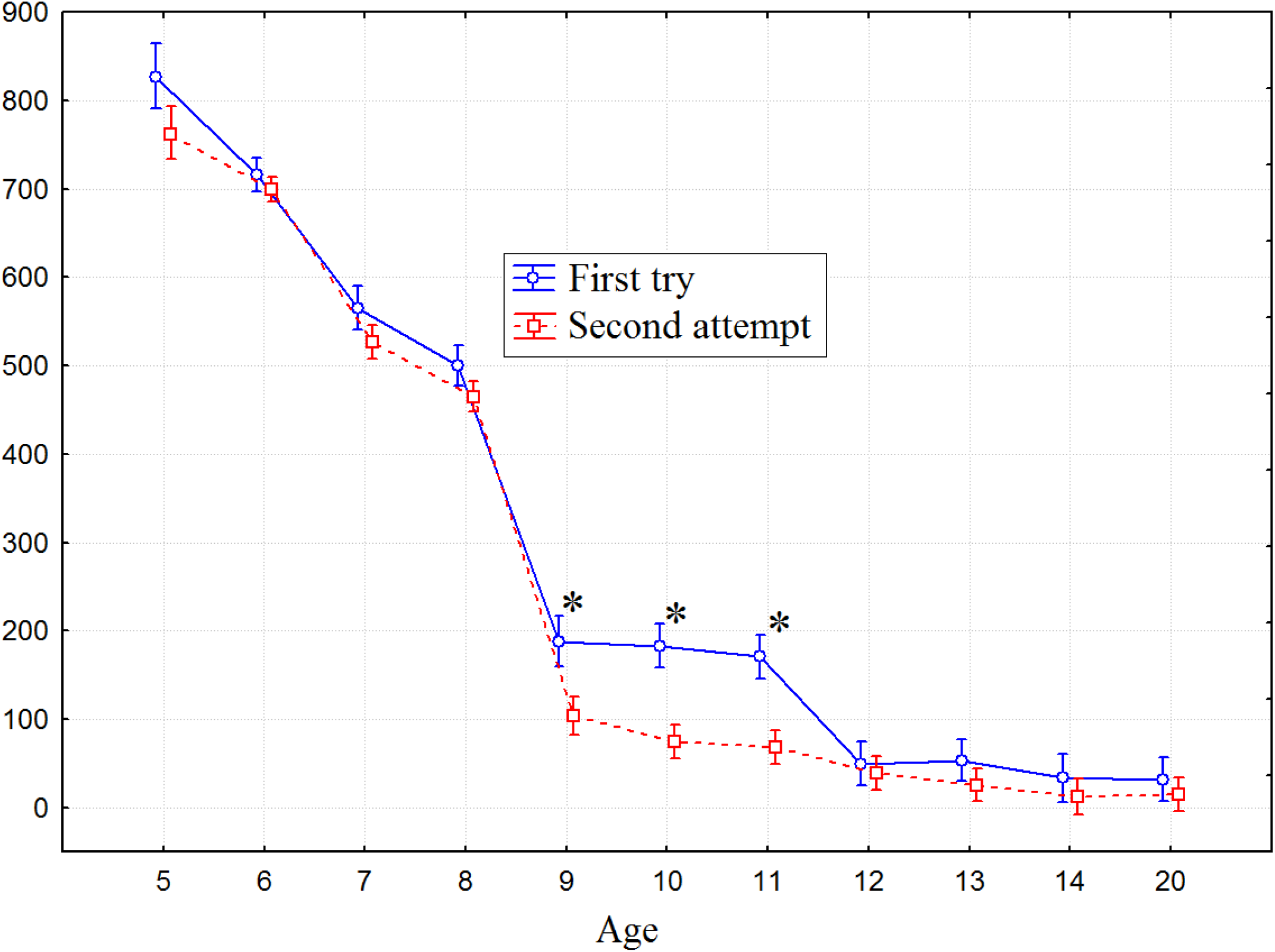
The difference between anticipation and real time of conturing (|antisipation of time – real time|). Gray color – first try, black – second attempt.

Children 9-11 years old demonstrated significant improvement of accuracy of time anticipation after the prompt (F(10, 302)=5,8404, p=,00000) compared to other aging groups (Figure 3). The error of anticipation of event’s duration at the first was the highest in 5 children, decreased during maturation and reached a plateau at the age of 12: the significant differences between children over 12 years old and adults were not detected. On the second attempt the plateau was reached at the age of ten.

## IV. Discussion

Our findings supported that a child’s concept of time and ability to evaluate, estimate, and plan the time develops during maturation (Piaget, 1981; Smythe, 1957). At the same time each component of time perception had its own critical period or zone of proximal development (Sargent, 2014). In particular, we found that the critical period of ability to estimate event’s duration was about seven years old and the zone of proximal development for ability to anticipate of event’s duration was from nine to eleven years old. The results mean that these abilities were actively developing during these ages and children were sensitive to perceive information related with the timing ability. Children from 5 to 8 years old overestimated time of Schulte tables’ reading, however unlike to 5 years old children of 6 to 8 years old could improve their inaccurate estimation after receiving a prompt. Nine years old children showed the accuracy of time estimation similar to adults and receiving a prompt didn’t improve their already sufficiently accurate estimation. This result indicated that age from 6 to 8 years old could be a zone of proximal development for task of estimation of event’s duration.

Another important finding of the present study concerns the dependence of children’s timing ability on both the difficulty of the task and their level of cognitive development. Numerous studies have shown that an individual’s time sensibility reaches an adult-like level at about 8–9 years of age, although some differences in time judgment persist in relation to more difficult tasks until the teenage years (Droit-Volet, 2013; Zélanti & Droit-Volet, 2011). A similar trend regarding the development of time perception was found in our research. More specifically, it was found that the ability to estimate ten-second and one-minute intervals develops earlier than the ability to plan and estimate the time of one’s own activity. Thus, the development of timing abilities, especially in the case of explicit time perception, stems from multiple factors, including experience of the temporal regularities of events, awareness of the passing of time, and development of general cognitive capacities (Droit-Volet, 2016; Droit-Volet & Zélanti, 2013). Moreover, we identified a number of neuropsychological indicators that, regardless of age, were found to influence a child’s timing abilities. In particular, the accuracy of the assessment of time was found to be significantly lower among those children who achieved worse results for the attention and memory tests. Hence, the awareness of time depends on a child’s attention and short-term memory capacities, which are related to the maturation of both the prefrontal cortex and the hippocampus (Wittmann, 2007). Previous findings also demonstrated that preschoolers had difficulty perceiving explicit and abstract representation of time, this ability of their duration judgements became better as cognitive and executive function develops with maturation of the prefrontal cortex (Coull et al., 2018).

Our data corresponded to previous findings showed that children between the ages of 6 and 10 years were characterized by intensive development of proactive control which allowed children to anticipate and prepare for upcoming events and required high level of working memory performance (Chevalier, 2014). Children under 8 years old mainly used reactive control activated only when the event had already occurred and had difficulties to plan the upcoming event (Pani, 2013). At the age of 8, children were able to use both types of control depending on the situation and only from 10 year old children preferred to use proactive control where possible (Chevalier et al, 2015). Proactive control required for the anticipation of event’s duration was provided by a wide neural network including prefrontal cortex (Zandbelt, 2013) which kept maturated during childhood and reached a peak in the cortical thickness by 11 years. The detected mistakes during dynamic praxis tests correlated in our study with the error of the time anticipation could be associated with the development of proactive control. Summarizing our findings we hypothesized that development of proactive control and other higher mental functions associated with the activity of prefrontal cortex contribute to zone of proximal development for the ability of events’ anticipation (Chevalier et al, 2013).

The relationship between the accuracy of an individual’s perception of time and that individual’s attention level has previously been reported. A number of researchers have demonstrated that young children’s poor judgments during the completion of classical Piagetian tasks are not due to their inability to correctly judge time, but rather to their limited attentional capacities (Droit-Volet, 2006). Other authors have suggested that attention-deficit hyperactivity disorder (ADHD) could be caused by a distorted sense of time; in which time passes so quickly that concentration becomes difficult (Goddard, 2000). Children and adolescents with ADHD display impairments in terms of both their duration discrimination and the precision with which they reproduce the intervals during the estimation task (Toplak, 2003). Further, children who exhibit an attention deficit have been found to make shorter reproductions of 15- and 60-second intervals (Barkley, 2001). Additionally, the visual-spatial memory has been found to be a significant predictor of difficulties in visual and auditory duration discrimination tasks at intervals 1000 ms in an ADHD sample (Toplak, 2005).

In the present study, certain other timing abilities, such as an improvement in the assessment of time, were found to be more pronounced in those participants who made a lower number of spatial and quasi-spatial verbal mistakes. Such data support previous assertions that explicit time perception depends on the level of an individual’s general cognitive capacities (Droit-Volet & Zélanti, 2013), which are also complemented by other studies that have reported an inverse relationship between the brain processing speed and the perceived passage of time (Goddard, 2000). At the same time, the development of time judgment was closely related to the development of spatial cognition (Coull et al., 2018), associated with the maturation of temporal, parietal and frontal areas (Nagy, 2004).

## V. CONCLUSION

The findings showed that the timing abilities are related with the general development of cognitive functions: the accuracy of the assessment of time was significantly lower in those children who achieved worse results for the attention and memory tests; the improvement of the assessment of time in relation to reading a Schulte table correlates with a higher level of spatial and verbal thinking. Zone of proximal development of ability to anticipate or plan the events duration was found in children from 9 to 11 years old.

## CONFLICT OF INTEREST

The authors declare that the research was conducted in the absence of any commercial or financial relationship that could be construed as a potential conflict of interest.

## FUNDING

This study was funded by the Russian Science Foundation (Grant No. 20-68-46042).

## Notes

### Competing Interest Statement

The authors have declared no competing interest.

